# Distribution of intraperitoneally administered D_2_O in AQP4-knockout mouse brain after MCA occlusion

**DOI:** 10.1101/2022.10.02.510567

**Authors:** Takuya Urushihata, Hiroyuki Takuwa, Manami Takahashi, Je◻ Kershaw, Sayaka Shibata, Nobuhiro Nitta, Yasuhiko Tachibana, Masato Yasui, Makoto Higuchi, Obata Takayuki

## Abstract

As the movement of water in the brain is known to be involved in neural activity and various brain pathologies, the ability to assess water dynamics in the brain will be important for the understanding of brain function and the diagnosis and treatment of brain diseases. Aquaporin-4 (AQP4) is a membrane channel protein that is highly expressed in brain astrocytes and is important for the movement of water molecules in the brain. In this study, we investigated the contribution of AQP4 to brain water dynamics by administering deuterium-labeled water (D_2_O) intraperitoneally to wild-type and AQP4 knockout (AQP4-ko) mice that had undergone surgical occlusion of the middle cerebral artery. Water dynamics in the infarct region and on either side of the anterior cerebral artery (ACA) was monitored with proton-density-weighted imaging (PDWI) performed on a 7T animal MRI. D_2_O caused a negative signal change quickly after administration. The AQP4-ko mice showed a delay of the time-to-minimum in both contralateral and ipsilateral ACA region compared to wild-type mice. Also, only the AQP4-ko mice showed a delay of the time-to-minimum in the ipsilateral ACA region compared to the contralateral side. In only the wild-type mice, the signal minimum in the ipsilateral ACA region was higher than that in the contralateral ACA region. In the infarct region, the signal attenuation was slower for the AQP4-ko mice in comparison to the wild-type mice. These results suggest that AQP4 loss affects water dynamics in the ACA region not only in the infarct region. Dynamic PDWI after D_2_O administration may be a useful tool for showing the effects of AQP4 in vivo.

**Highlights:**

- The measurement of brain water dynamics with D_2_O.
- *In vivo* investigation of AQP4 loss and focal brain ischemia.
- AQP4 loss affects water dynamics in the ACA region not only in the infarct region.

## 1. Introduction

Water dynamics in the brain has recently been a topic attracting significant interest. Some reports have suggested that water dynamics affects sleep level and dementia (Rasmussen et al., 2018; Taoka and Naganawa, 2020). In previous studies, two-photon microscopy imaging (Iliff et al., 2012) and MRI with an ^17^O isotope tracer (Kudo et al., 2018) have been used for real-time monitoring of in vivo brain water dynamics. However, as ^17^O isotope tracers are very expensive, there is a need for a non-invasive, safe and inexpensive method that can be used to assess and diagnose patients with brain diseases.

Aquaporin is a protein complex embedded in cell membranes that facilitates the exchange of water molecules from one side of the membrane to the other (Ohene et al., 2019; Pavlin et al., 2017; Vajda et al., 2004; Zhang et al., 2019). One subtype, aquaporin-4 (AQP4), is known to express in the end feet of astrocytes. It regulates the flow of water from intravascular and extracellular compartments into the intracellular compartment, and is involved in the removal of excess water in the brain (Badaut et al., 2007b; Rasmussen et al., 2018; Verkman, 2009). AQP4 is known to be involved in many brain diseases such as stroke and dementia (Abe et al., 2020; Aoki et al., 2003; Manley et al., 2000; Xu et al., 2015; Zeppenfeld et al., 2017), and AQP4 knockout animals are often used to investigate the function of AQP4. AQP4 knockout (AQP4-ko) mice could be useful for clarifying the role of AQP4 in brain water dynamics and for developing new treatments for brain diseases that target AQP4.

In direct measurements using an optical imaging technique called coherent anti-Stokes Raman scattering (CARS) imaging, and indirect studies using di usion MRI, di◻erent levels of AQP4 expression have been found to a ect cell membrane water permeability (Obata et al., 2018). CARS is sensitive to H_2_O, but is insensitive to deuterium-labeled water (D_2_O) signal. Therefore, CARS can be used to observe intracellular and extracellular water exchange after rapid replacement of H_2_O to D_2_O in the extracellular space. Applying the same procedure during MRI may enable the measurement of water dynamics *in vivo*. Similar to the case of CARS, it is expected that the process of exchanging H_2_O with D_2_O will correspond to signal loss in proton-density-weighted imaging (PDWI).

In the early stages of a mouse model of focal cerebral ischemia, it has been reported that the diffusion-weighted imaging (DWI) signal change in AQP4-ko mice is di erent from that for wild-type mice (Urushihata et al., 2021). Water movement to astrocytes via AQP4 is an important process in the early stages of ischemia-induced brain edema formation. AQP4 expression is also sharply increased in ischemic brain edema (De Castro Ribeiro et al., 2006; Hirt et al., 2009), and AQP4-ko or inhibitor administration has been shown to be effective in reducing cellular edema (Akdemir et al., 2014; Igarashi et al., 2011; Manley et al., 2000; Yao et al., 2015). Therefore, we have assessed the early stages of focal cerebral ischemia in a mouse model to highlight the effects of AQP4 on water dynamics. In this study, dynamic PDWI was performed after intraperitoneal administration of D_2_O to compare the distribution of water in wild-type and AQP4-ko mice brains after middle cerebral artery occlusion (MCAO).

## 2. Materials and Methods

### 2.1. Animal preparation

A total of five C57BL/6J wild-type mice (both male and female, 20-30 g, 8-10 weeks; Japan SLC, Hamamatsu, Japan), and five AQP4-ko mice (both male and female, 20-30 g, 8 to 10 weeks) generated as described previously (Ikeshima-Kataoka et al., 2013; Kato et al., 2014) (acc. no. CDB0758 K: http://www.cdb.riken.jp/arg/mutant%20mice%20list.html), were used in the MRI experiments. All mice were housed individually in separate cages with water and food ad libitum. The cages were kept at a temperature of 25◻°C in a 12-h light/dark cycle. Overall, no clear differences in body weight and size were observed for any of the mice. MCAO was performed for all animals at 3 hours before beginning the dynamic PDWI scans. The occlusion was performed using the Tamura method (Tamura et al., 1981), where a permanent occlusion is made at the proximal branch of the MCA in the left cerebral cortex. In this animal model, ischemia in the MCA region occurs soon after MCAO, and the infarction expands and peaks at 24 hours after surgery (Brint et al., 1988; Garcia et al., 1993; McCullough and Liu, 2011; Tamura et al., 1981). All experiments were performed in accordance with the institutional guidelines on humane care and use of laboratory animals, and were approved by the Institutional Committee for Animal Experimentation of the National Institutes for Quantum and Radiological Science and Technology (QST).

### 2.2. Magnetic resonance imaging measurements

MRI measurements were performed with a 7T animal MRI (Bruker Biospin, Ettlingen, Germany). The mice were initially anesthetized with 3.0% isoflurane (Escain, Mylan Japan, Tokyo, Japan), and then with 1.5% to 2.0% isoflurane and a 1:5 oxygen/room-air mixture during the MRI experiments. Rectal temperature was continuously monitored with an optical fiber thermometer (FOT-L, FISO, Quebec, QC, Canada), and maintained at 37.0±0.5°C using a heating water pad. Warm air was provided with a homemade automatic heating system regulated by an electric temperature controller (E5CN, Omron, Kyoto, Japan) throughout all experiments. During MRI scanning, the mice lay in a prone position on a MRI-compatible cradle and were held in place with handmade ear bars.

At the beginning of the scanning session, DWI with a four-shot pulsed-gradient spin-echo (PGSE) EPI sequence (TR◻=◻3◻s, TE◻=◻115◻ms, matrix size◻=◻128◻×◻128, spatial resolution◻=◻0.02◻×◻0.02 mm^2^, slice thickness◻=◻1.5◻mm, gradient directions = 3, b = 1000 s/mm^2^, Δ = 100 ms, δ = 7 ms) was taken to confirm ischemic changes. Dynamic PDWI was performed using a four-shot spin-echo EPI sequence (TR = 5s, TE = 11ms, FOV = 19.2mm, matrix size◻=◻64×64, slice thickness◻=◻1mm) to acquire an image every 20 s for 30 min (total 90 scans). Two minutes after scanning began, 1 ml of 99.8% D_2_O saline was administered intraperitoneally (Fig. 1). One problem for the administration of D_2_O to mice is the very small blood volume of a mouse (2-3 ml). The circulation system would be severely a ected (Kselíková et al., 2019) if an intravenous injection of 1ml D_2_O was performed. To avoid this problem, the D_2_O was administered intraperitoneally so that the exchange between D_2_O and intravascular H_2_O is less dramatic.

**Figure. 1.**
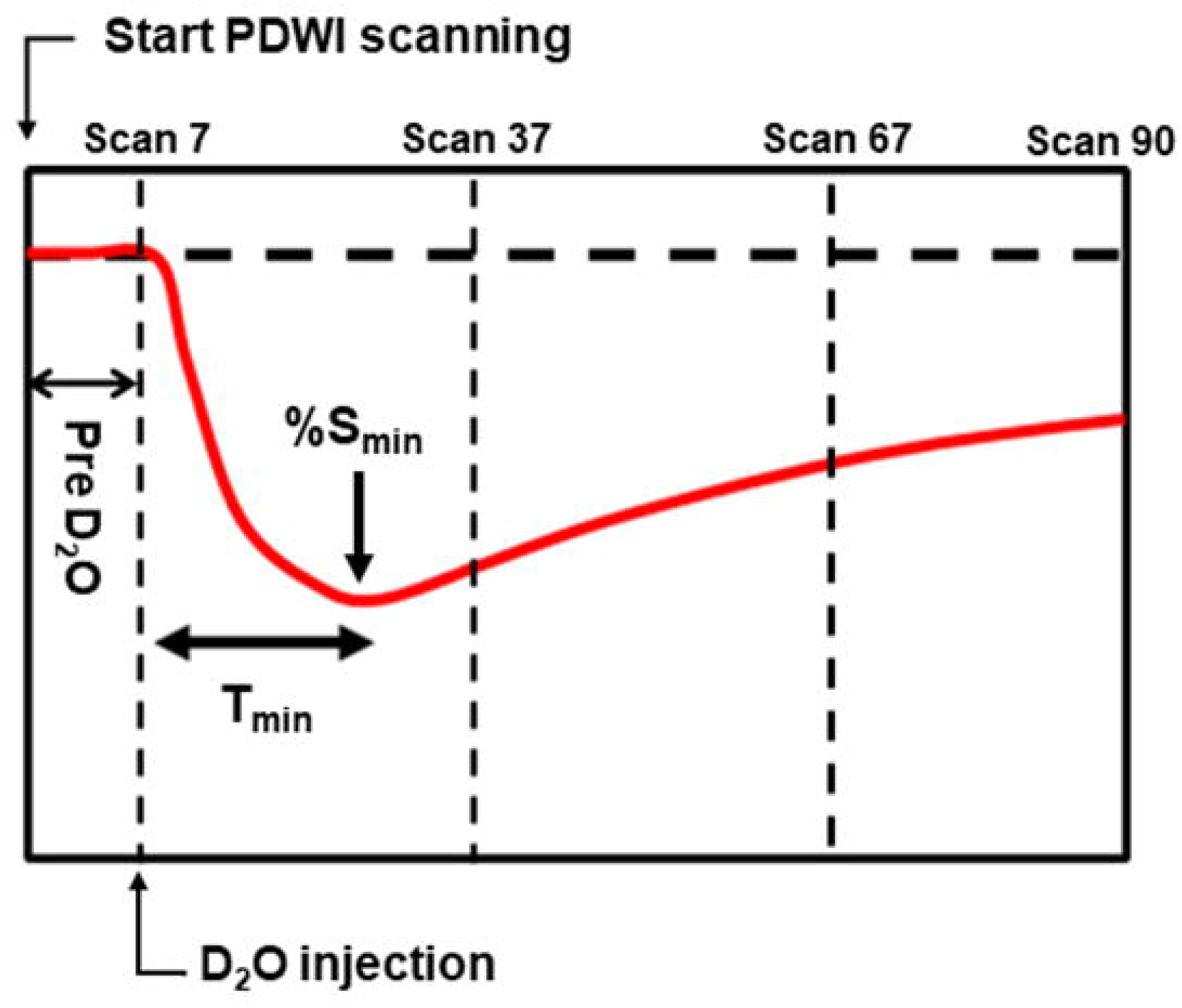
Schematic defining the timing of the D_2_O administration in comparison to the PDWI. PDWI images were acquired every 20 s for 30 min (90 scans total). Two min after scanning started, D_2_O was administered intraperitoneally. *T_min_* indicates the time from administration to the signal minimum.

### 2.3. PDWI data processing

#### 2.3.1 Signal normalization for maps and time-intensity curves

Analysis of the PDWI data was performed in MATLAB, version R2019a (MathWorks, Natick, MA, USA). After discarding the first scan, the next five scans acquired in the first two minutes prior to D_2_O administration were averaged and used as the reference image S_0_ for the pre-D_2_O signal. For all scans thereafter the percent signal change, *%S(t)*, was calculated as,

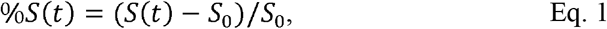

where *S(t)* is the signal intensity at time *t* (min).

#### 2.3.2 ROI analysis

In order to evaluate PDWI signal changes after D_2_O administration, regions-of-interest (ROIs) were drawn at the following three locations: high signal region in the MCA area on the DWI (Infarct), the cortex in the anterior cerebral artery (ACA) territory on the same side as the MCAO (Ipsi-ACA), and the cortex in the opposite ACA territory (Contra-ACA) (Fig. 2A). The mean time-dependent signal in each ROI between scan 7 and scan 37 (Fig. 1) after D_2_O administration was interpolated with an 8th order polynomial. The time from D_2_O administration to the signal minimum (*%S_min_*) was defined as the time-to-minimum (*T_min_*). Since the transfer rate of D_2_O from the peritoneal cavity to the blood is different for different individuals, the following standardization of the signal was performed.

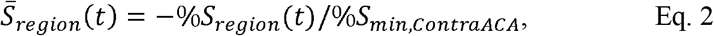

where “region” corresponds to one of the ROIs (i.e. Infarct, Ipsi-ACA, or Contra-ACA), and %*S_min, ContraACA_* is the minimum of *%S(t)* for the Contra-ACA ROI. After this procedure, the average value before D_2_O administration is zero for all ROIs, and the minimum value of the Contra-ACA signal 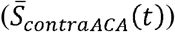 is −1.

**Figure. 2.**
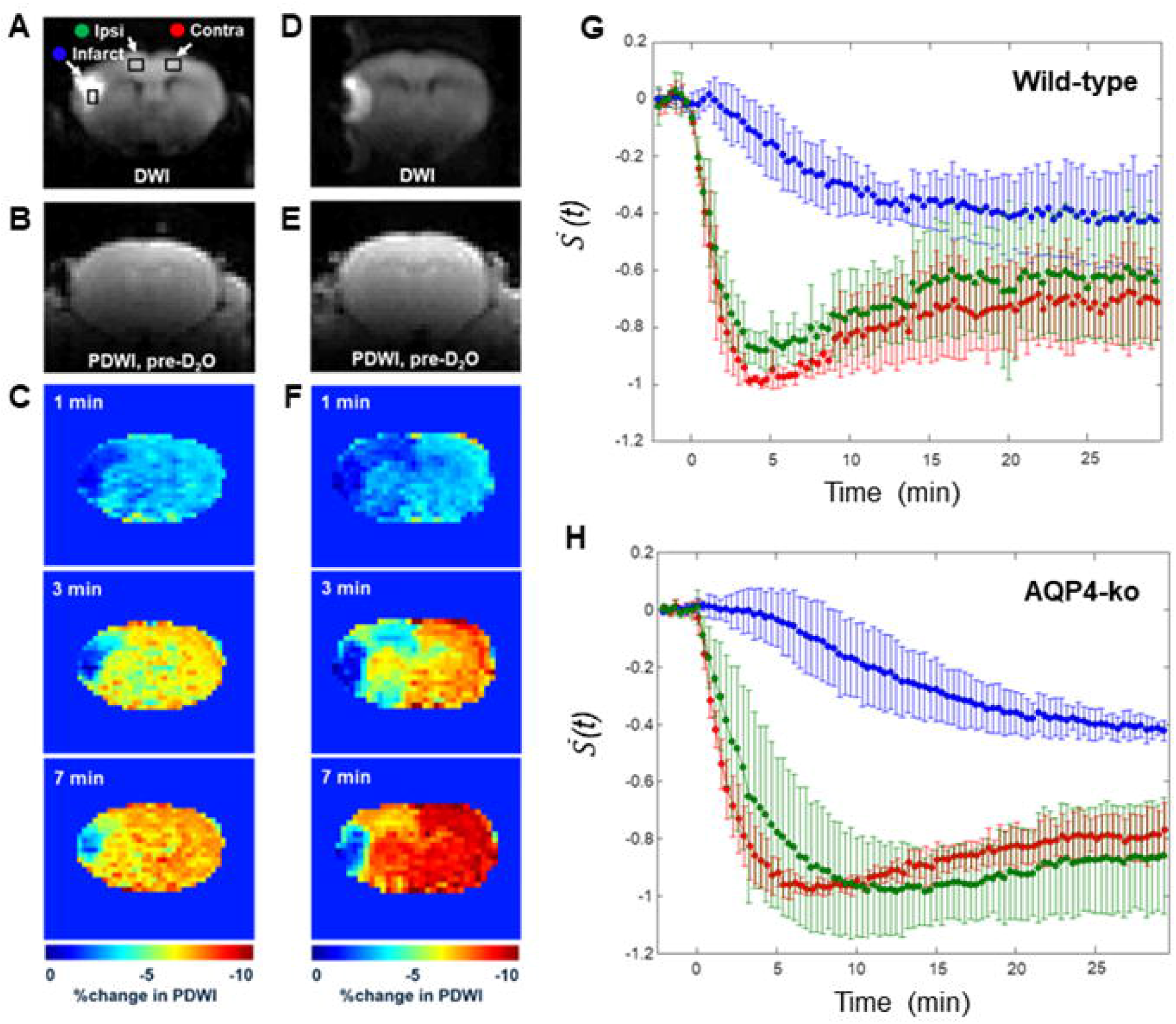
Water dynamics measurements after intraperitoneal administration of deuterium-labeled water. (A-C) Representative DWI images, Pre-D_2_O baseline PDWI images and % signal change at 1, 3 and 7 min after D_2_O administration for the wild-type mice. (D-F) A similar se t of images for the AQP4-ko mice. (G) Standardized signal changes in each ROI for the wild-type mice. (H) Standardized signal changes in each ROI for the AQP4-ko mice. Error bars indicate standard deviations across animals. The Infarct (blue), Ipsi-ACA (green), and Contra-ACA (red) ROIs are shown on the DWI image in (A).

For the Infarct ROI, a signal minimum was never reached for 8 of the 10 subjects so, as an alternative method to characterize signal change, the slope α of the time-intensity curve between 3 and 5 minutes after D_2_O (3-5min) was estimated.

### 2.4. Statistical analysis

Statistical analyses were performed with the Statistics and Machine Learning Toolbox of MatLab. All results are presented as mean◻±◻standard deviation over animals. Student’s unpaired t-test was used to compare *T_min_* and α between mouse genotypes, while the paired t-test was used to compare between regions. The normality of each data set was confirmed with the Kolmogorov-Smirnov Test. A p-value < 0.05 was interpreted as being statistically significant.

## 3. Results

### 3.1 Dynamic PDWI after intraperitoneal D2O injection

In the DWI performed prior to dynamic PDWI, all mice showed a high signal region in the ipsilateral MCA territory (infarct region in Figs. 2A and D). PDWI signal decreases in the brain were observed for all mice immediately after intraperitoneal administration (Figs. 2B-C and E-F), which reflects the transfer of D_2_O into the brain. For the wild-type mice, signal change in the Ipsi-ACA ROI was similar to that observed in the Contra-ACA ROI (Fig. 2G), whereas for the AQP4-ko mice the signal from the Ipsi-ACA ROI appears to be slightly delayed in comparison to that from the Contra-ACA ROI (Fig. 2H). The signal from the Infarct ROI was substantially different in comparison to the other ROIs for both mouse types (Figs. 2G and H).

### 3.2 Time-to-minimum (*T_min_*) and signal change in the Infarct ROI

Figure 3A shows that there was no di◻erence in *T_min_* between the Contra-ACA ROIs and Ipsi-ACA for the wild-type mice, while for the AQP4-ko mice a significant di erence was observed. *T_min_* between mouse type was also compared and there were also significant di erences in the estimates for the Contra-ACA and Ipsi-ACA ROIs, although the significance was quite weak for the Contra-ACA ROI (p=0.028).

**Figure. 3.**
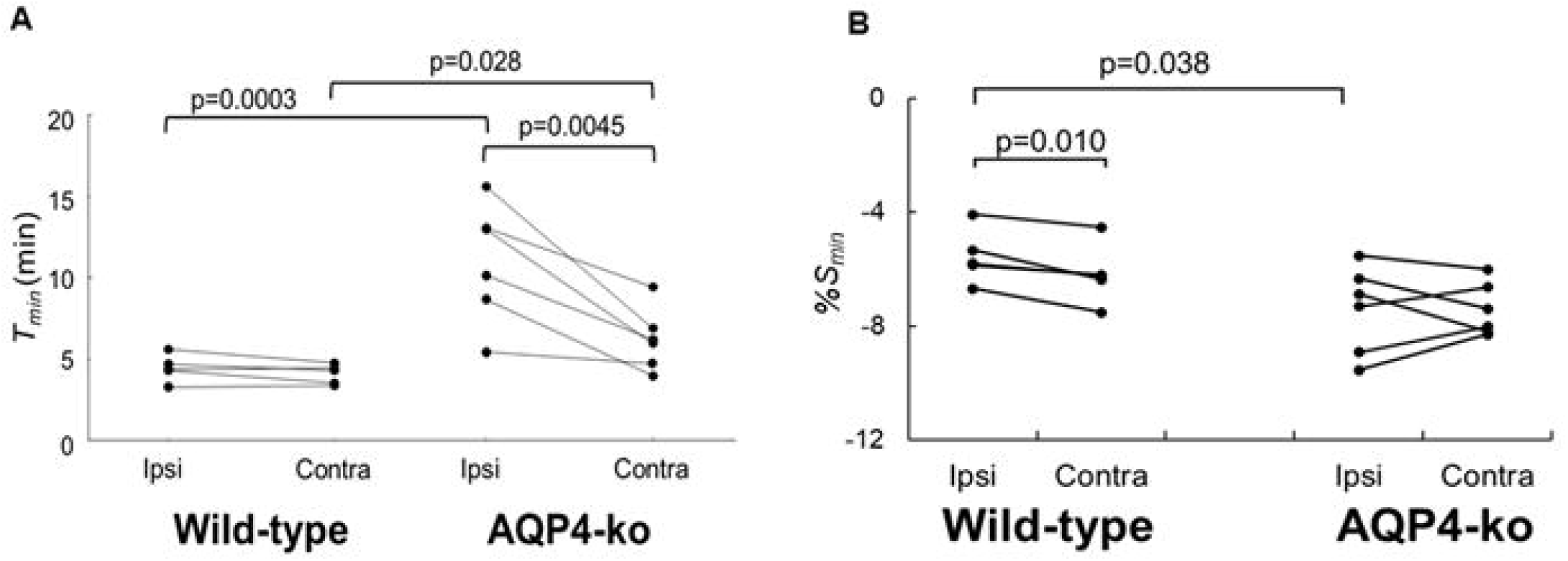
Time-to-min (*T_min_*) and signal minimum (*%S_min_*) in the ACA ROIs after D_2_O administration. (A) *T_min_* for the wild-type (n=5) and AQP4-ko (n=5) mice. (B) *%S_min_* for both mouse genotypes. Results for the same animal are connected by lines. Comparisons between genotypes were made using an unpaired t-test, while the comparison between the ipsi and contralateral sides were made with a paired t-test. The resulting p values are shown at the top of the figure.

The *%S_min_* was significantly higher in the Ipsi-ACA ROI than in the Contra-ACA ROI for the wild-type mice (Fig. 3B). There was no significant difference in the corresponding comparison for the AQP4-ko mice. When comparing the *%S_min_* between genotypes, the Ipsi-ACA ROI had a significantly higher value for the wild-type mice.

Since the data did not reach a minimum in the Infarct ROI for 8 out of the 10 mice, the slope α of the signal attenuation at 3-5 minutes after D_2_O administration was used to evaluate signal change (Fig. 4A). For the wild-type mice α was −0.016 ± 0.0070/min, which was smaller than the α of −0.0030 ± 0.0069/min for the AQP4-ko mice. This difference was significant (p=0.019; Fig. 4B).

**Figure. 4.**
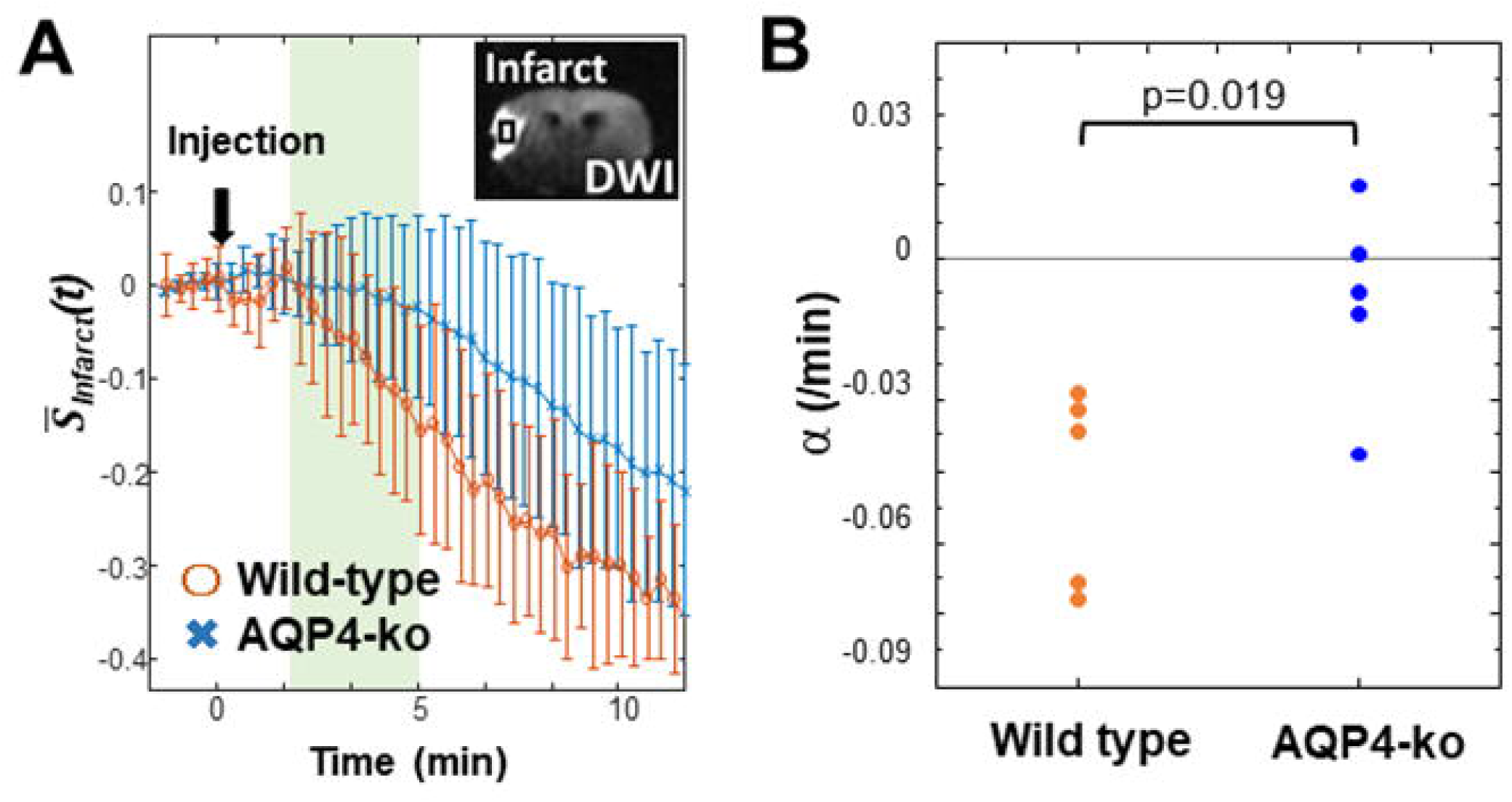
Signal attenuation in the Infarct region. (A) Standardized PDWI signal in the Infarct region of the wild-type (orange) and AQP4-ko (blue) mice. The slope α is calculated from the signal recorded at 3-5 min after D_2_O injection. (B) The estimates of α for the two groups were compared using an unpaired t-test and the p value is shown at the top of the figure.

## 4. Discussion

In this experiment, we used dynamic PDWI after a intraperitoneal D_2_O injection to observe the biodistribution in brains with MCA occlusion. Our method depicted the AQP4-expression-related temporal and spatial differences in water dynamics associated with focal cerebral ischemia. Significant differences in the PDWI signal change were observed between wild-type and AQP4-ko mice.

Deuterium is a stable isotope of hydrogen, and D_2_O is non-toxic unless in high volumes (Kselíková et al., 2019). Since D_2_O is undetectable in ^1^H MRI, its mixing with H_2_O in vivo causes concentration-dependent attenuation of the PDWI signal. So D_2_O is effectively a negative contrast agent. The motion of D_2_O in vivo is the same as H_2_O so it can pass through aquaporins (Obata et al., 2018), making it a potential means to assess aquaporin function and perfusion. D_2_O was administered intraperitoneally in the present study because of the possibility that intravenous administration of a high volume of D_2_O might affect the cardiovascular system (Kselíková et al., 2019). Comparison of intraperitoneal and intravenous administration of Evans blue to evaluate blood brain barrier (BBB) permeability has been reported to show no difference in brain accumulation at any time point (Manaenko et al., 2011). The same is probably true for D_2_O, which is easily passes through the BBB. Intraperitoneal administration was suitable for D_2_O imaging because it causes less damage to the body and allows experiments to be repeated.

### Time-to-minimum (*T_min_*) and Signal-at-min (*%S_min_*)

There was no significant difference in *T_min_* between the ipsilateral and contralateral sides for the wild-type mice, while for the AQP4-ko mice the value was significantly larger on the ipsi side (Fig. 3). This means that changes to the signal in the Ipsi-ACA ROI of the AQP-ko mice was delayed. In general, when ligation is performed at the main trunk of the MCA, arterial blood is partially compensated by flow from the ACA, which means that supply to the ACA region is reduced (Dziennis et al., 2015). The elongation of *T_min_* for the Ipsi-ACA ROI of the AQP4-ko mice seems to be naturally a ected by this CBF reduction. The fact that there was no significant change for the wild-type mice is curious. It is possible that the number of active AQP4 sites increases in the Ipsi-ACA region of the wild-type mice so that the water transfer to brain is maintained even under the condition of low CBF (Badaut et al., 2007a).

In *T_min_*, there was a significant difference between wild-type and AQP4-ko even in Contra-ACA. This may be due to the delayed flow of water from the capillary to the perivascular space (Mestre et al., 2020) and the enlargement of the extracellular spece, as suggested by diffusion MRI studies (Urushihata et al., 2021).

The *%S_min_* in the Ipsi-ACA ROIs of the wild-type mice was higher than that in the Contra-ACA ROIs for all animals (Fig. 3B). As there was no difference in *T_min_* for these ROIs, the concentration of D_2_O in the Ipsi-ACA area was about 12% less than that in the Contra-ACA region. This suggests that dispersion of the D_2_O is slightly reduced due to the MCA occlusion even in the wild-type mice. On the other hand, there was no difference between the *%S_min_* of the Ipsi-ACA ROIs and Contra-ACA ROIs for the AQP4-ko mice (Fig. 3B). This may be due to the signal reduction by D_2_O influx to the enlarged extracellular mentioned in *T_min_*, which may cancel the *%S_min_* change as seen in the wild-type mice. The wild-type mice had a significantly higher *%S_min_* than the AQP4-ko mice for the Ipsi-ACA ROI, but the reason for this is unclear.

### Signal change in the Infarct area

Signal attenuation in the infarct area tended to be slower for the AQP4-ko mice. The infarct region is thought to have almost no blood flow for both the wild-type and AQP4-ko mice due to the MCA occlusion. Since AQP4 expression is known to increase rapidly after occlusion in the core and border regions of the lesion (De Castro Ribeiro et al., 2006; Hirt et al., 2009), the di erence in signal changes in the Infarct ROI may be due to the faster inflow of D_2_O from the surrounding intact area in the wild-type mice. This may be due to the larger influx of water from surrounding tissue to the infarct lesion via the perivascular space in the wild-type mice (Mestre et al., 2020).

### Conclusion

Dynamic PDWI after administration of D_2_O is useful for investigating the dynamics of D_2_O in the brain, and is thought to be effective in showing the effects of AQP4. Under severe ischemic conditions, the differences between the wild-type and AQP4-ko mice were significant. This may be because some compensation mechanism for intracerebral water transfer fails for AQP4-ko mice in a pathological state.

## Abbreviations

ACA: Anterior cerebral artery
AQP4: Aquaporin-4
BBB: Blood brain barrier
CARS: Coherent anti-Stokes Raman scattering
D_2_O: Deuterium-labeled water
DWI: Diffusion-weighted imaging
MCAO: Middle cerebral artery occlusion
MRI: Magnetic resonance imaging
PDWI: Proton-density-weighted imaging
ROI: Regions-of-interest

## 7. Acknowledgements

The authors thank Mr. Takeharu Minamihisamatsu for help with animal management. This work was supported by a Grant-in-Aid for Scientific Research (KAKENHI, #15H04910 [TO]) from the Japan Society for the Promotion of Science (JSPS).

